# Epidemiological evaluation and identification of the insect vector of soybean stay-green associated virus

**DOI:** 10.1101/2023.03.13.532384

**Authors:** Ruixiang Cheng, Rong Yan, Ruoxin Mei, Yaodi Wang, Wei Niu, Hao Ai, Sijing Qiao, Mengjia Xu, Wei Yu, Wenwu Ye, Yuanchao Wang, Xiaorong Tao, Xueping Zhou, Yi Xu

## Abstract

In recent years, the emergence of soybean stay-green syndrome (SGS), also referred to as “zhengqing”, in the Huang-Huai-Hai region of China has resulted in significant yield losses, with some areas experiencing a complete reduction in seed yield. SGS is a phenomenon characterized by the delayed senescence of soybean, resulting in stay-green leaves, flat pods, and stunted seed development at harvest. Our group was the first to identify a distinct geminivirus, named soybean stay-green associated geminivirus (SoSGV), as the causative agent of SGS by fulfilling Koch’s postulate. To further understand the epidemiology of SoSGV, in this study, we collected 368 stay-green samples from 17 regions in 8 provinces including the Huang-Huai-Hai region and surrounding areas of China. The results showed that 228 samples tested positive for SoSGV (61.96%), and 96.93% of these positive samples showed severe pod deflation. Our epidemiological assessment reveals SGS caused by the SoSGV is prevalent in the fields, and it is undergoing geographical expansion and genetic differentiation. Additionally, we determined the other natural hosts grown in the Huang-Huai-Hai region of China. By capturing insects in the field and conducting laboratory vector transmission tests, we confirmed that the common brown leafhopper (*Orosius orientalis*) is the transmitting vector of SoSGV. With a better understanding of the epidemiology of SoSGV and its transmission, we can develop more effective strategies for managing and mitigating its impact on soybean yields.

## 1. Introduction

Soybean is a highly valuable legume crop worldwide, providing 25% of the global edible oil supply and two-thirds of the global concentrated protein for livestock feed (https://www.fas.usda.gov/data/oilseeds-world-markets-and-trade). However, the emergence of soybean stay-green syndrome (SGS), also known as “zhengqing”, in the Huang-Huai-Hai region of China in recent years has caused significant yield losses, with some areas experiencing a complete loss of seed yield [1-3]. SGS is known as delayed senescence at harvest, which is characterized by the retention of green leaves, poor pod production, and stunted seed development at harvest [1-4]. Recently, our group discovered for the first time that a new distinct geminivirus named soybean stay-green associated geminivirus (SoSGV) is the etiology of soybean stay-green syndrome by fulfilling Koch’s postulate [2]. The SoSGV infection alone can cause soybean stay-green symptoms, including delayed leaf senescence (stay-green), increased numbers of abnormal seeds, and many flat pods. This novel distinct virus is a monopartite single-strand DNA virus, which is most likely formed by intergeneric recombination of geminiviruses [2].

In our previous study, we discovered that SoSGV not only infects soybean, but also various experimental hosts, such as *Nicotiana benthamiana, N. tabacum, N. glutinosa*, and *Datura stramonium*. However, the natural hosts of SoSGV, particularly crops grown in the same area as soybean in the Huang-Huai-Hai region, were unknown. Additionally, we have previously investigated the transmission mode of SoSGV and provided compelling evidence that it cannot be transmitted through mechanical inoculation or seed transmission. Although the virus could reach the seed coat of soybean, it was unable to penetrate the embryo and cotyledon, indicating that seed transmission of SoSGV is unlikely. We further confirmed that the predominant *Bemisia tabaci* Mediterranean (Q biotype) in the Huang-Huai-Hai region does not act as a vector for the transmission of SoSGV. Based on the finding that the CP structure of SoSGV was highly similar to that of CP from *Mastrevirus*, we proposed that SoSGV might be transmitted by leafhoppers [2].

Otherwise, the mechanisms behind the formation of soybean stay-green syndrome are not well understood, but it is believed to be caused by the disruption of the interaction between source activity and sink capacity due to external factors [5-7]. Previous studies have shown pod removal and seed injury, as well as feeding by bean bugs *Riptortus pedestris* (Fabricius) (*Hemiptera: Alydidae*) and red-banded stink bugs *Piezodorus guildinii* (Westwood) (*Hemiptera: Pentatomidae*), can cause delayed senescence, probably by impairing the sink capacity [7, 8]. To gain a better understanding of the epidemiology of SoSGV and identify the main factors contributing to SGS in the Huang-Huai-Hai region, we conducted a study in 2022 to collect and test stay-green soybean plants at harvest from 17 different regions spanning 8 provinces. Our epidemiological assessment of SoSGV reveals it is undergoing geographical expansion and genetic differentiation. We also identified the natural hosts and transmitting vector of SoSGV. With a better understanding of the epidemiology of SoSGV and its transmission, we can develop more effective strategies for managing and mitigating its impact on soybean yields. Our study provides valuable information for guiding agricultural production and integrated control measures for SoSGV, which can help reduce yield losses and ensure sustainable soybean production in China.

## 2. Materials and Methods

### 2.1 Field Survey and Sampling

Whole plant samples showing delayed senescence were collected during the harvest period from September to November 2022 from 17 distinct geographic regions across 8 provinces in China (Figure 1a). Samples were collected objectively based on plants remaining green, without relying on virus symptoms. Overall, 368 samples were collected in these 17 regions and all samples (weeds and pea) were grouped into lots of three, respectively prior to virus testing. Details of the collection site and numbers of collected samples are summarized (Table 1). Samples were stored frozen at −80 °C until subsequent DNA extraction and PCR testing.

**Figure 1.**
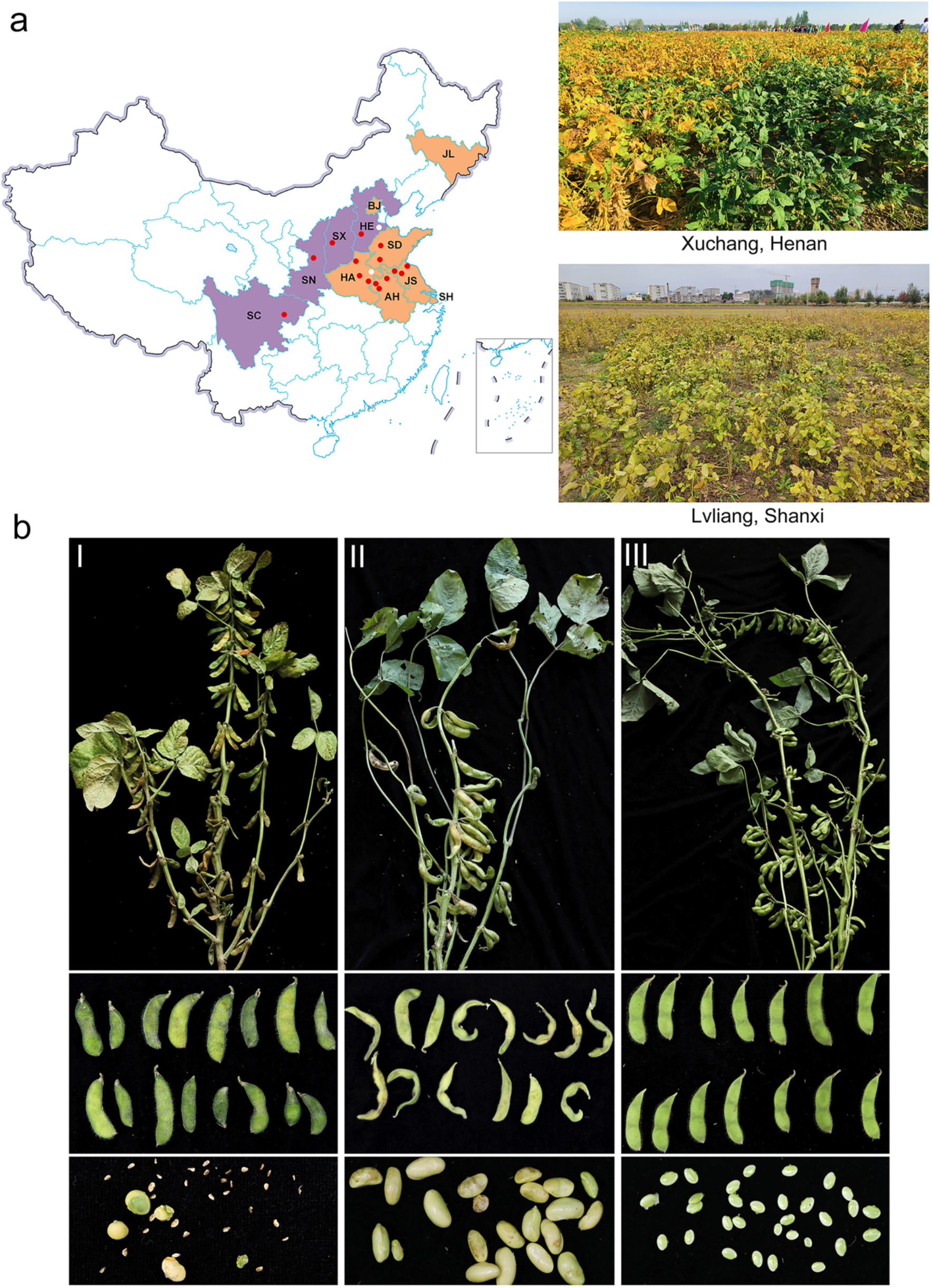
Distribution and symptoms of SoSGV. (a) A geographic map highlighting all regions with confirmed incidences of SoSGV (in orange), as well as regions with new emerging cases reported in the current study (in purple). Red dots represent sites where SoSGV was detected, while white dots indicate no SoSGV was found in the samples. The right panel of (a) shows the field symptoms commonly associated with SoSGV infection. Two regions displaying these symptoms are shown. (b) Symptom classification of collected stay-green soybean samples. (I), Class I refers to samples with obvious flat pod symptoms and a normal number of pods, while Class II (II) indicates samples with non-flattened pods and a significantly reduced number of pods. Class III (III) denotes samples with non-flattened pods and a normal number of pods.

**Table 1.**
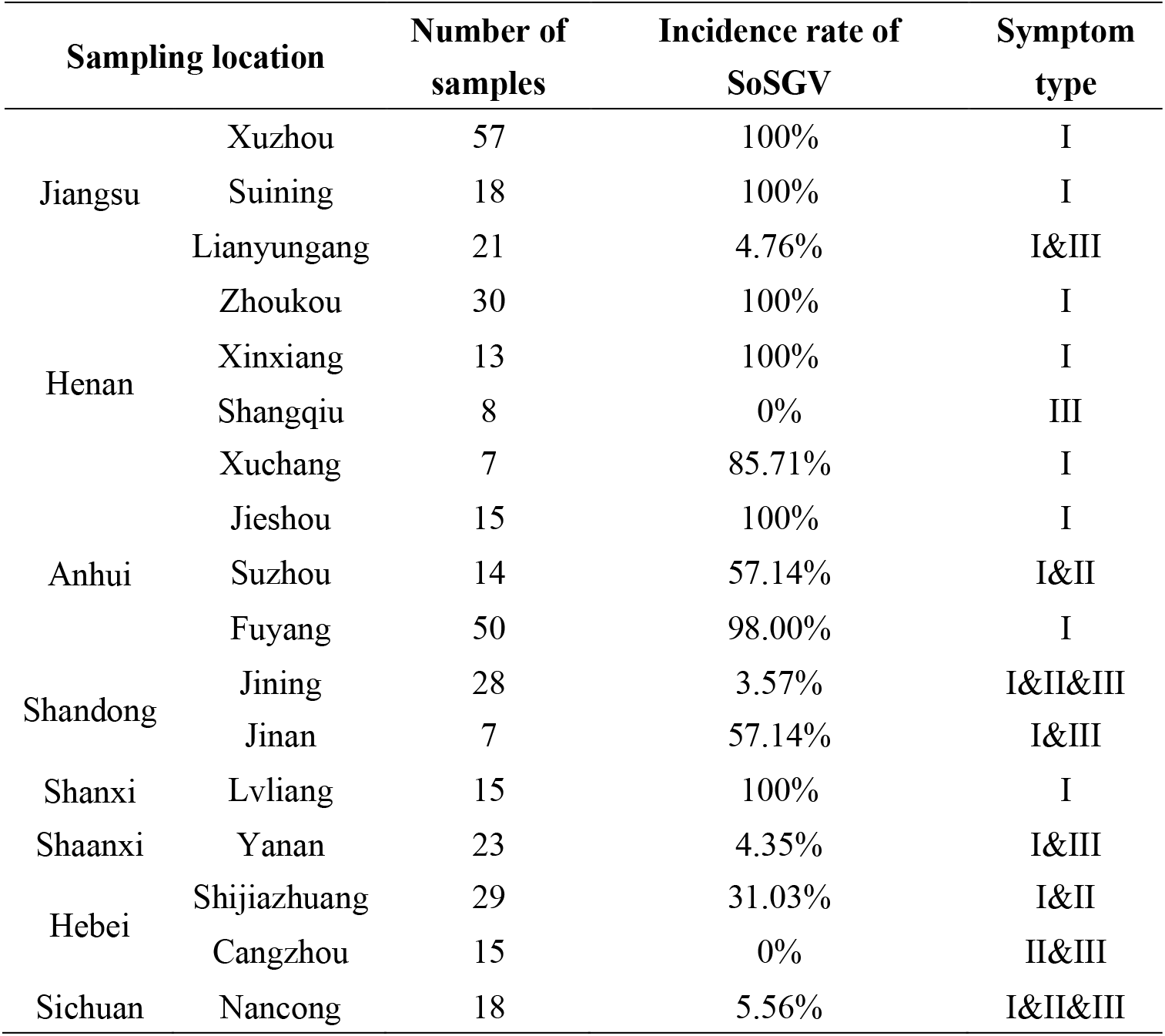
PCR detection results of soybean samples and classification of diseased soybean types

### 2.2. DNA extraction and PCR

The total DNA of plant samples was extracted with CTAB method-based extraction procedure [9]. To extract DNA from a single insect, we placed it in a PCR tube and added lysis buffer consisting of 10 mM tris-HCl, 0.45% NP40, 50 mM KCl, 0.45% Tween-20, 0.2% gelatin, 60 mg/L protease K, with pH adjusted to 8.4. Next, we added 6-8 glass beads with a diameter of 1 mm and ground the mixture on a grinder at 60 Hz for 240 seconds. After rapid centrifugation, we incubated the sample in a 65-degree water bath for 60 minutes. To inactivate the protease, we boiled the sample in water for 10 minutes. Finally, we collected the supernatant by rapid centrifugation and used it for PCR detection or stored it at -20 °C.

PCR was conducted with the specific primer designed for CP (CP-F: 5’-ATGGATTACAGCAGGAAGAGG-3’; CP-R: 5’-TTACAATTTGCTCTTGAAATACGT-3’) to detect the presence of SoSGV in each sample. The full-length SoSGV genome of were amplified using a pair of back-to-back primers, FL-F (5’-CGCTGTTAAGCGCCTTGGCGTAAGC-3’) and FL-R (5’-CGGCAGTAAGTCAGAGGCTCTTAA-3’). For cloning the partial *COI* gene of the leafhopper, a pair of degenerate primer was designed based a previous study with minor modification [10], and COI-F (5’-AACTTTATACTTTATmTTTGGTATTTGATCAGG-3’) and COI-R (5’-AAwACTGGTAGTGACAryAATArTAG-3’). All the PCR products were separated using 1% agarose gel, and gel extraction and sequencing were subsequently performed. The *Orosius orientalis* partial CDS of *COI* gene cloned from leafhopper collected from Fuyang has been deposited to NCBI with GenBank submissions number OQ472322.

### 2.3. Agrobacteria and agroinfiltration

Peas, chickpeas, cowpeas, and broad beans are owned by our lab. Potato is a laboratory-owned Desiree variety. As for corns, all experimental cultivars including ‘B73’, ‘Suyu Nuo’, ‘Suyu 29’, ‘Run Yangyu’ and ‘Suyu 39’ were provided by Dr. Qin Gu from Nanjing Agricultural University. And Chinese milk vetch and lucerne (*Medicago sativa* L.) were kindly provided from Dr. Bo Yang from Nanjing Agricultural University. All the plants were grown in a greenhouse at 26-28 °C and 65% relative humidity under 16/8 h day/night conditions. For infectivity, the plasmids containing SoSGV infectious clone were transformed into *A. tumefaciens* strain EHA105 by electroporation method. Then agrobacteria harbored SoSGV infectious clone was suspended with buffer (10 mM MES, 10 mM MgCl2, 200 μM acetosyringone) for 2 hours at OD_600_=1.0, then infiltrated into plant leaves with 4-6 true leaves. To further confirm the infectivity, *A. rhizogenes* strain K599-mediated hairy root transformation inoculation method was carried out on corns simultaneously as described in our previous paper [2]. Agrobacterium transformed with pBINPLUS empty vector were infiltrated into plant as negative control.

### 2.4. Phylogenetic analysis

Using the samples collected from Huang-Huai-Hai region in China, full-length of 44 SoSGV sequences were cloned and sequenced. Whole genome sequences of representative members of each of the fifteen regions that SoSGV was present. Sequences were aligned by ClustalW implemented in MEGA software version X [11]. and phylogenetic trees were constructed by the Maximum Likelihood method [11]. One thousand replications were analyzed to enhance robustness. To identify the taxonomy of leafhopper, we designed primers targeting the mitochondrial cytochrome oxidase subunit 1 (COI) gene of the *Orosius* genus and performed sequence analyses and integrative taxonomy based on a previous study [10]. Then a phylogenetic tree was also constructed by the ML method with 1,000 replications for each bootstrap values using MEGA X.

### 2.5. Insect capture and transmission assay

Insects with sucking or piercing-sucking mouthparts including aphids, leafhoppers, bean bugs, and whiteflies were captured from a soybean field in Fuyang, Anhui province where all soybean samples from this field were tested positive for SoSGV (data not shown). A portion of each insect samples collected was stored at −20 °C for DNA extraction and for SoSGV testing by PCR and the remaining was cage-reared along with soybean plants collected from the same field sesame plants under greenhouse conditions with 26-28 °C and 65% relative humidity under 16/8 h day/night conditions. Insect samples were identified to species by morphological characteristics using a stereomicroscope. Three SoSGV-viruferious adult aphids, whiteflies, leafhoppers, or bean bugs were then transferred to each caged healthy soybean (4-5 leaf stage), respectively. The insects were allowed a 48-hour access period to transmit the virus. After inoculation, the plants were treated with an insecticide and placed in an insect-proof greenhouse. Four weeks later, we tested the plants for infection using PCR. Those that tested positive were kept in the greenhouse, monitored for the appearance of symptoms for up to three months. Two cultivars including *Glycine max* cv. ‘Williams 82’ and cv. ‘Zhonghuang 13’ were used.

## 3. Results

### 3.1 SoSGV is widely distributed in soybean fields and continues to expand its range to other areas

To advance our understanding of the epidemiology of SoSGV, we conducted a study from September to November 2022 in which we collected and tested 368 soybean plants that remained green at harvest from 17 distinct geographic regions across 8 provinces in China for the presence of the virus (Figure 1a). The results of the study revealed that SoSGV was present in 61.96% (228 out of 368) of the stay-green soybean samples analyzed (Table 1). Specifically, 100% of the samples collected from Xuzhou in Jiangsu Province, Zhoukou in Henan Province, Xinxiang in Henan Province, Jieshou in Anhui Province, and Lvliang in Shanxi Province tested positive for SoSGV. The incidence of SoSGV in samples from Xuchang in Henan Province, Suzhou in Anhui Province, Fuyang in Anhui Province, and Jinan in Shandong Province was also notably high, with more than 50% of the samples testing positive. In contrast, the incidence of SoSGV in samples from Lianyungang in Jiangsu Province, Jining in Shandong Province, Yan’an in Shaanxi Province, and Nanchong in Sichuan Province was found to be below 10%. No SoSGV was detected in samples collected from Shangqiu in Henan Province or Cangzhou in Hebei Province. The statistical analysis results are presented in Table 1.

Furthermore, the study revealed that the symptoms of SGS exhibited variability among the stay-green soybean samples tested (Figure 1b). Based on the number of pods and the degree of pod filling observed in the samples, the symptoms were classified into three types as depicted in Figure 1b: Type I, characterized by evident flat pod and normal pod number; Type II, featuring no pod collapse but a reduced number of pods; and Type III, displaying no flat pod and a normal pod number. Of the 368 samples collected, 227 samples displayed symptoms characteristic of Type I, while 82 samples exhibited symptoms of Type II and 59 samples of Type III. In terms of the virus-carrying rate, 97.36% of the Type I samples were tested positive for SoSGV, with a 6.1% virus-carrying rate in Type II and a 3.39% rate in Type III (Table 1). These results indicate that infection with SoSGV resulted in abnormal filling and flattening of soybean pods, which is consistent with the previously defined symptoms of SGS. Notably, samples exhibiting symptoms of Type II and Type III did not exhibit flattening of pods (Figure 1b). In addition to its previously identified presence in the Huang-Huai-Hai Region, the spread of SoSGV has now been discovered in numerous other soybean producing regions, including Hebei, Shaanxi, Shanxi, and Sichuan Provinces, based on the results of our large-scale survey (Table 1 and Figure 1a).

### 3.2 Phylogenetic analysis of SoSGV in stay-green soybean samples

To better understand the spread and evolution of SoSGV, a phylogenetic analysis was conducted on a collection of soybean plant samples collected from various regions. Specifically, 44 samples of 228 stay-green soybeans from 15 locations in 8 provinces were selected for SoSGV full-length cloning and sequencing. The genomic sequences of the SoSGV isolates were compared, revealing 92.08-99.9% nt identity among them (Supplementary File S1). Subsequent phylogenetic analysis of the complete genomes of 44 SoSGV isolates demonstrated that the Lvliang and Shijiazhuang isolates form a separate, coherent, and bootstrap-supported clade (Figure 2). In addition to geographical expansion, our epidemiological assessment of SoSGV in China suggests that it is undergoing genetic differentiation.

**Figure 2.**
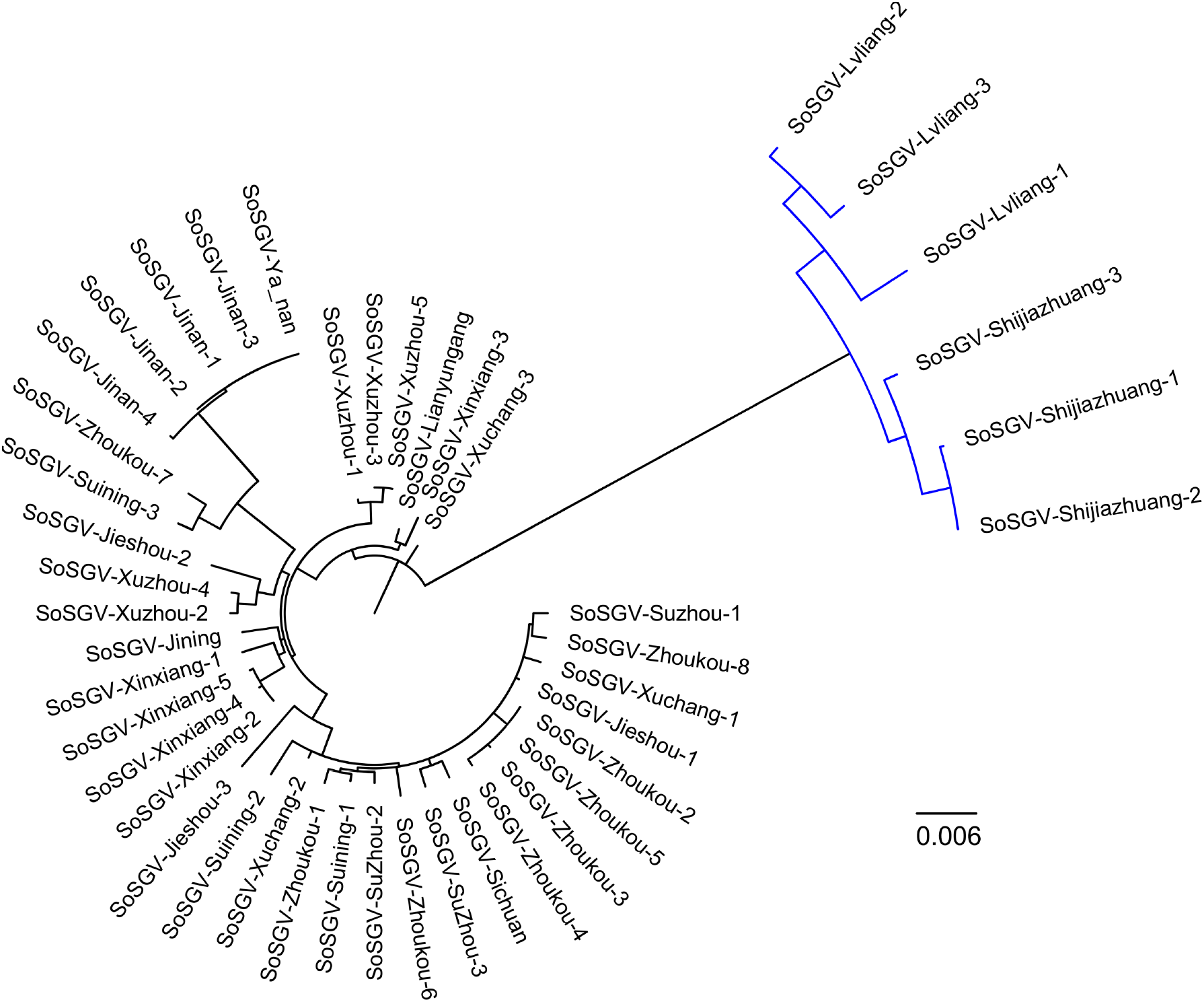
Phylogenetic tree based on forty-four full genome sequences of SoSGV from 15 regions in 8 provinces. The tree was constructed using the maximum-likelihood (ML) method in the MEGA X software. The bootstrap confidence values generated by 1000 replications and separate clades were marked with blue color.

### 3.3 Determination of the natural host range of SoSGV

In our previous study, we discovered that SoSGV can infect not only soybean, but also other experimental hosts such as *N. benthamiana, N. tabacum, N. glutinosa*, and *D. stramonium* [2]. To expand our understanding of the host range of SoSGV, particularly in crops that are grown in the same area as soybean in the Huang-Huai-Hai region, we conducted further tests using the infectious clone of SoSGV. The results, presented in Tables 2 and Figure 3, showed that SoSGV can also infect peas, chickpeas, and Chinese milk vetch in the legume family. Among these crops, pea is a widely cultivated crop globally, and Chinese milk vetch is valued for its role as a green manure and feed for livestock. SoSGV infection in peas resulted in a stay-green phenotype similar to that seen in soybean (Figure 3a). In contrast, Chinese milk vetch infection caused the leaves to noticeably shrivel (Figure 3b). Chickpea plants infected with SoSGV exhibited weaker growth compared to healthy plants (Figure 3c). Notably, we found that SoSGV was unable to infect potatoes, a member of the *Solanaceae* family (Table 2). Furthermore, we tested five corn cultivars, a gramineous crop grown alongside soybean, and found that SoSGV was unable to infect corn (Table 2).

**Table 2.**
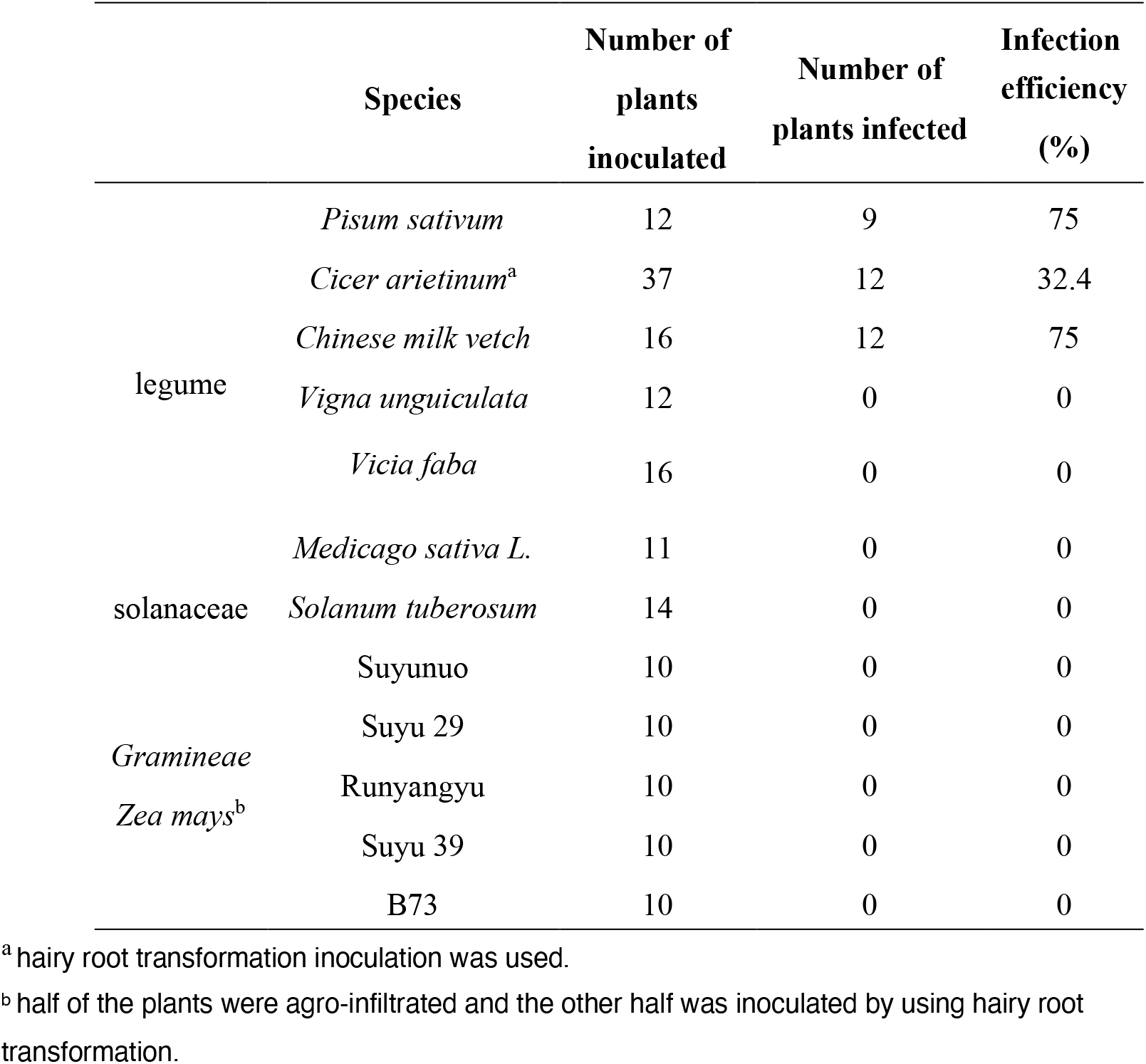
Determination of the natural hosts of SoSGV

**Figure 3.**
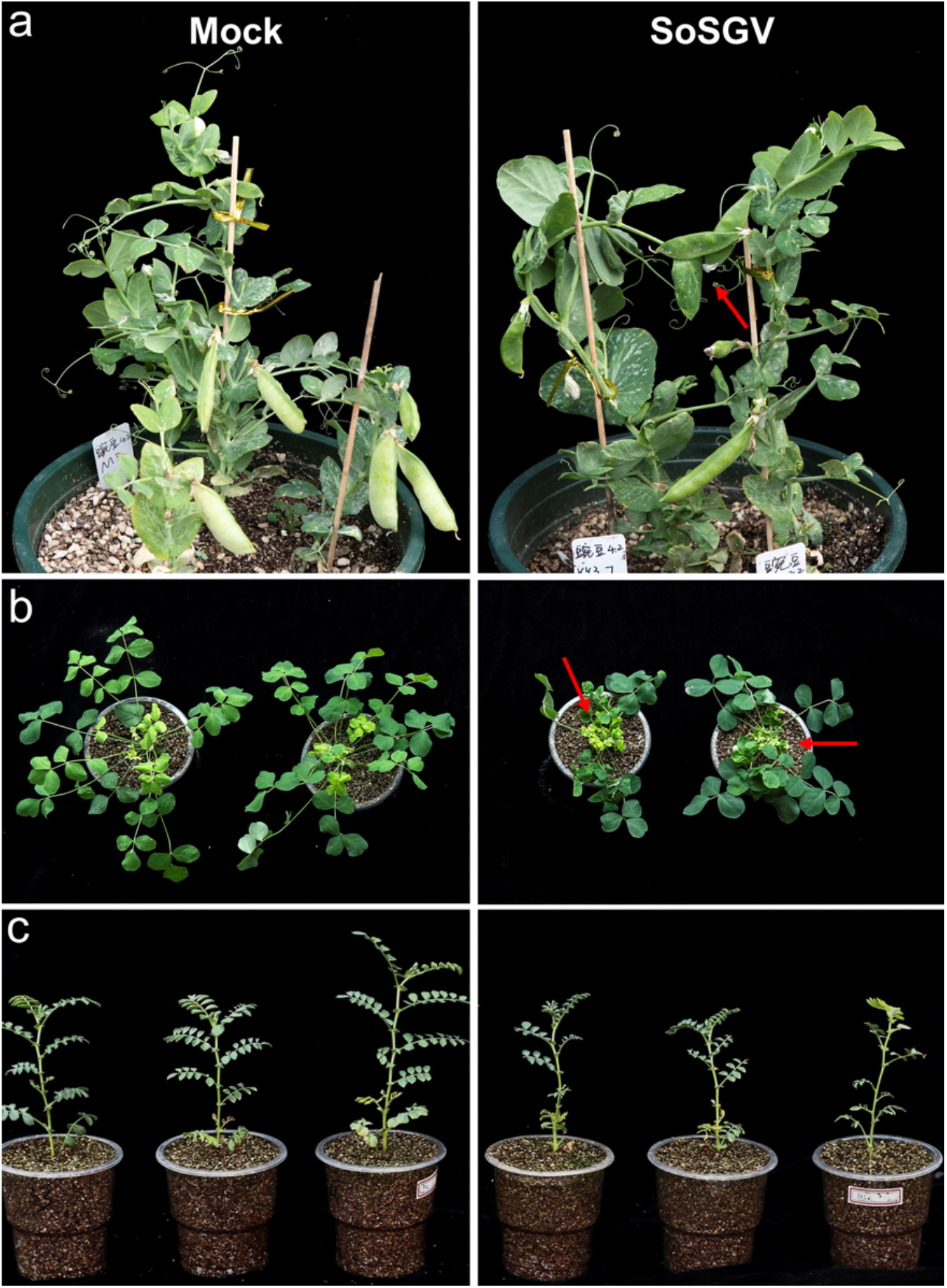
Symptoms associated with SoSGV infection of peas, chickpeas, and Chinese milk vetch. Peas (a), chickpeas (b), and Chinese milk vetch (c) were inoculated with SoSGV infectious clone at 3 weeks post sowing, respectively. The symptoms were photographed at 35 days post inoculation of SoSGV. Red arrows in A indicated clear flat pod symptom and in B, noticeably shriveled leaves are observed, also marked by red arrows.

### 3.4 Identification of insect vector transmitting SoSGV

To determine the insect(s) responsible for the transmission of SoSGV, we captured insects with sucking or piercing-sucking mouthparts, such as aphids, leafhoppers, bean bugs, and whiteflies, from a soybean field in Fuyang, Anhui Province, where all soybean samples collected from this field were tested positive for SoSGV (data not shown). To further confirm the transmission vector(s), we placed 15 healthy soybean plants in cages containing insects feeding on SoSGV-infected soybeans. Four weeks after infestation, all 15 plants were tested for the presence of SoSGV, and the results showed that they were infected, indicating that the insects containing the vector(s) transmitting the virus (Table 3). We then conducted laboratory transmission tests using each insect species to identify which could transmit SoSGV. In each assay, three insects were transferred to a healthy soybean plant, and the inoculation access period was set at 2 days. Four weeks later, leaf samples were collected and tested for the presence of SoSGV. The results showed that only leafhoppers were able to transmit SoSGV to soybean, while all other insects failed to do so (Table 3). The tests were conducted using two different soybean cultivars, and the results were consistent. After three months of transmission by leafhoppers, the infected plants exhibited several symptoms that are characteristic of stay-green symptoms, including thicker and darker-green foliage, numerous flat pods, and seeds with abnormal pod-filling (Figure 4).

**Table 3.** Determination of the insect vector of SoSGV

| Insect species | No. of plant infected with SoSGV/no. of plant tested |  |
| --- | --- | --- |
|  | Williams 82 | Zhonghuang 13 |
| Mixed insects | 15/15 | 15/15 |
| Bean bug | 0/15 | 0/15 |
| Whiteflies | 0/15 | 0/15 |
| Aphid | 0/10 | 0/10 |
| Leafhopper | 10/10 | 10/10 |

**Figure 4.**
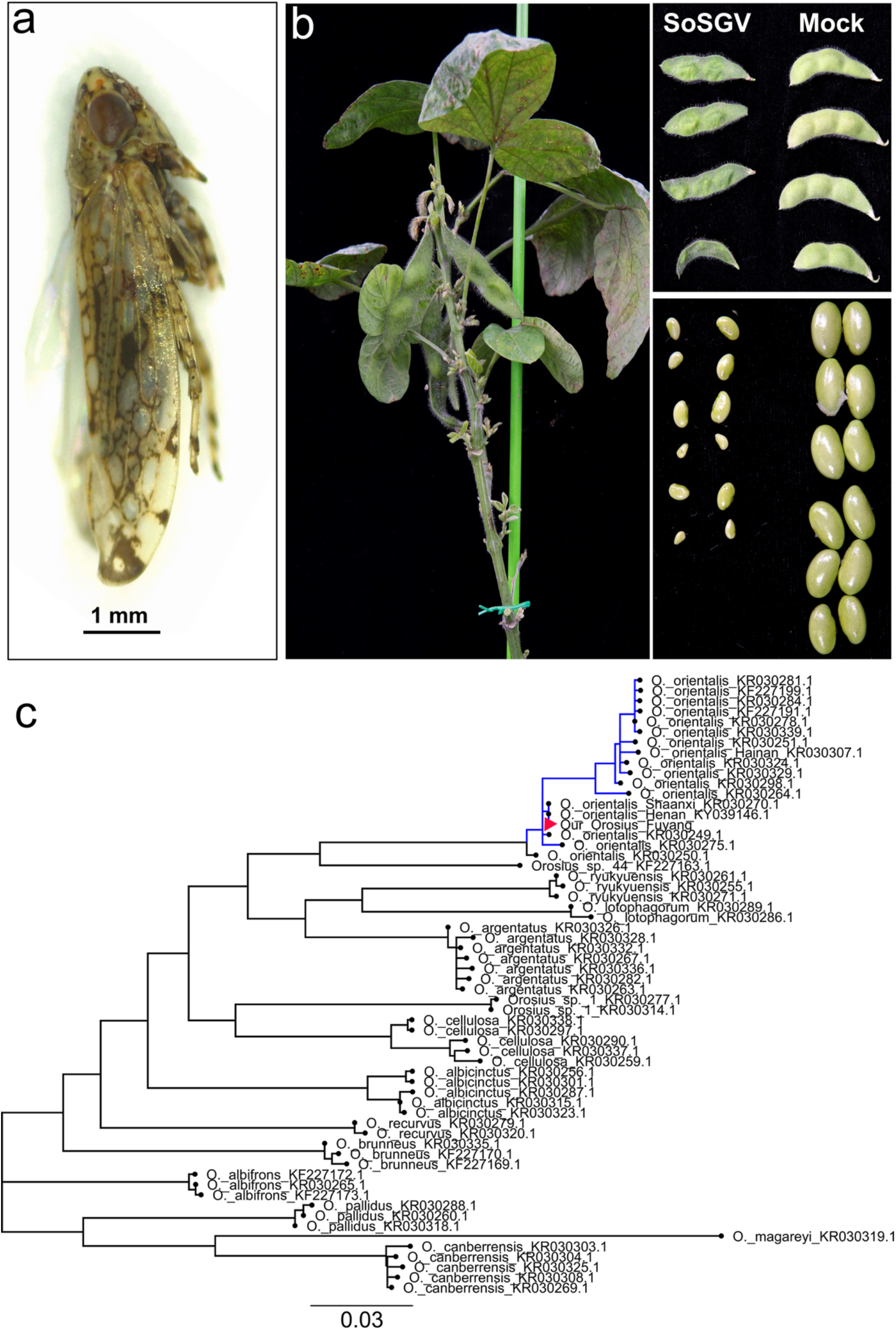
Identification of *O. orientalis* as the transmission vector of SoSGV. (a) Habitus picture of an adult common brown leafhopper (*O. orientalis*) collected in the field. (b) Soybean plants that were fed on by SoSGV-viruliferous *O. orientalis* developed thicker and darker-green foliage, numerous flat pods, and seeds with abnormal pod-filling at 3 months post infestation. (c) COI DNA barcoding sequences of 58 *Orosius* reference species including all 12 known *Orosius* species and two indeterminate *Orosius* (sp. # 1; sp. # 44) were retrieved from NCBI (https://www.ncbi.nlm.nih.gov). Phylogenetic tree constructed by the ML method showing relationships among our *Orosius* and reference *Orosius* species. The bootstrap confidence values generated by 1000 replications. The GenBank number of all the reference sequences used are listed.

Compared to the green leafhoppers, *Cicadella viridis* (Linnaeus) and *Empoasca flavescens* (Fabricius), the leafhoppers we collected from the field exhibited distinctive brown morphological characteristics (Figure 4a). These leafhoppers are the second-largest insect population after whiteflies in the samples we collected. The adults were approximately 2.8 to 3.5 mm in length, with a pale-yellow head marked by an irregular dark-brown pattern, dark-brown eyes, and a pronotum displaying a pale yellow anterior third and a grey posterior two-thirds speckled with transverse dark-brown markings (Figure 4a). These morphological features suggested that the leafhoppers belonged to the *Orosius* genus. To confirm our hypothesis, we designed primers targeting the mitochondrial cytochrome oxidase subunit 1 (*COI*) gene of the *Orosius* genus, based on a previous study by Fletcher *et al*. [10]. We extracted DNA from ten individual adult leafhoppers, and their COI nucleotide sequences were found to be 100% identical. Therefore, we selected one sample for the construction of a phylogenetic tree, as shown in Figure 4c. Our leafhopper clustered with haplotypes of *O. orientalis* (Matsumura) (Homoptera: Cicadellidae), and was closely related to *O. orientalis* haplotypes collected from China [12]. Remarkably, the COI sequence of our *O. orientalis* (Fuyang) sample was 100% identical to that of a Henan haplotype (Genbank No. KY039146.1). This finding represents the first evidence that *O. orientalis* acts as the natural transmitting vector of SoSGV.

## 4. Discussion

In our previous study, we fulfilled Koch’s postulate and identified SoSGV as a causative agent of SGS for the first time. In this current study, we collected and analyzed 368 soybean stay-green samples from 17 regions in 8 provinces. Our results revealed that SoSGV was present in 61.96% of the samples, suggesting that it is the leading cause of soybean green stem syndrome (SGS) in the Huang-Huai-Hai region at present. Notably, samples were collected objectively based on plants remaining green, without relying on the presence of virus symptoms. During the harvest season, individual soybean plants with delayed leaf senescence that remained green were particularly noticeable. SoSGV was found to be sporadically distributed in the field, and it is spreading to other soybean producing areas (Figure 1a). Furthermore, SoSGV is still mutating and evolving, with isolates from Shijiazhuang and Lvliang forming distinct categories (Figure 2). In the future, we will be conducting a comparative analysis of the pathogenicity of different isolates, providing insights into the mechanisms that contribute to their virulence.

In addition, our study examined the natural host range of SoSGV by testing several crops that are commonly grown alongside soybeans. Our results showed that leguminous plants such as peas, chickpeas, and Chinese milk vetch can also be infected by SoSGV, in addition to soybeans (Table 2). In the Huang-Huai-Hai region, where peas are frequently planted during spring and early summer, their harvest season coincides with the soybean planting season, making peas a potential intermediate host of SoSGV. Chinese milk vetch, a commonly used green manure, was also found to be a potential host of SoSGV. By identifying the host range of SoSGV, our study sheds light on the occurrence pattern of SoSGV in the field, providing a scientific basis for more effective control of soybean stay-green disease.

*O. orientalis* (Matsumura) (Homoptera: Cicadellidae) is a significant vector of various viruses and phytoplasmas globally. It can transmit phytoplasmas that cause several economically important diseases, such as legume little leaf and tomato big bud [13], lucerne witches broom [14], potato purple top wilt [15], Australian lucerne yellows [16], and sesame phyllody. Additionally, *O. orientalis* is responsible for transmitting tobacco yellow dwarf virus (TYDV, genus *Mastrevirus*, family *Geminiviridae*) to beans, leading to bean summer death disease, and to tobacco, causing tobacco yellow dwarf disease [17]. Moreover, this leafhopper transmits Chickpea chlorotic dwarf geminivirus (genus *Mastrevirus*, family *Geminiviridae*), which is one of the viruses associated with chickpea stunt disease [18]. In this study, we demonstrated for the first time that *O. orientalis* is the natural transmitting vector of SoSGV. Notably, our previous research revealed that the CP structure of SoSGV is highly similar to that of CP from *Mastrevirus*, leading us to hypothesize that leafhoppers might transmit SoSGV [2]. Our experimental results align with this hypothesis. Although the common brown leafhopper is not native to China, it has been found in various locations worldwide, including mainland Australia and central Pacific islands, the Philippines, and Malaysia [10]. Further research is needed to investigate the distribution, host range, and potential impact on agricultural production in China.

## 5. Conclusions

In the present study, we conducted an overall survey that collected and tested stay-green soybean plants at harvest from 17 regions across 8 provinces of China. Our epidemiological investigation of SoSGV revealed that it is currently undergoing geographical expansion and genetic differentiation. Additionally, we were able to confirm the natural hosts of SoSGV, and for the first time, identified *O. orientalis* as the natural vector responsible for its transmission. With a better understanding of the epidemiology of SoSGV and its transmission, we can develop more effective strategies to manage and mitigate its impact on soybean yields.

## Supplementary Materials

The following supporting information can be downloaded at: https://www.mdpi.com, Supplementary File S1: Estimates of evolutionary divergence among 44 SoSGV isolates.

## Author Contributions

YX, XPZ, XRT, YCW conceived the project. RXC, RY, RXM, YDW, WN, HA, SJQ, MJX, WY and WWY performed the experiments. YX and RY wrote the manuscript. All authors read and approved the final manuscript.

## Funding

This work was supported by the National Key Research and Development Program of China (2022YFC2602000) and (2022YFF1001500), the Open Research Fund from State Key Laboratory for Biology of Plant Diseases and Insect Pests (SKLOF202205), and Jiangsu Agricultural Science and Technology Innovation Fund CX(22) 2039.

## Institutional Review Board Statement

Not applicable.

## Informed Consent Statement

Not applicable.

## Data Availability Statement

All the data used in this study are already provided in the manuscript in its required section. There is no underlying data available.

## Acknowledgements

We would like to express our great gratitude to China Agriculture Research System (CARS-004) for their assistance in collecting the soybean stay-green samples. These stations and institutes include Xuzhou, Zhengzhou, Shangqiu, Fuyang, Suzhou, Jining, Jinan, Fenyang, Yan’an, Cangzhou, Nancong, Zhoukou Academy of Agricultural Sciences, Anhui Academy of Agricultural Sciences and Hebei Academy of Agricultural Sciences. I am deeply grateful to my son Ethan for his tireless efforts in collecting leafhoppers from the fields.

## Competing interests

The authors declare that they have no competing interests.

## Notes

### Competing Interest Statement

The authors have declared no competing interest.

## References

1. Xu, C.; Han, T.; Wu, C., Discussion on the causes of staygreen syndrome for summer soybean and its preventive methods in the Huang-Huai-Hai region. Soybean Sci & Technol 2019, 160, (3), 22–28.

2. Cheng, R.; Mei, R.; Yan, R.; Chen, H.; Miao, D.; Cai, L.; Fan, J.; Li, G.; Xu, R.; Lu, W.; Gao, Y.; Ye, W.; Su, S.; Han, T.; Gai, J.; Wang, Y.; Tao, X.; Xu, Y., A new distinct geminivirus causes soybean stay-green disease. Mol Plant 2022, 15, (6), 927–930.

3. Wang, X.; Wang, M.; Wang, L.; Feng, H.; He, X.; Chang, S.; Wang, D.; Wang, L.; Yang, J.; An, G.; Wang, X.; Kong, L.; Geng, Z.; Wang, E., Whole-plant microbiome profiling reveals a novel geminivirus associated with soybean stay-green disease. Plant Biotechnol J 2022, 20, (11), 2159–2173.

4. Li, Q. L.; Zhang, Y. Y.; Lu, W. G.; Han, X. Y.; Yang, L. L.; Shi, Y. J.; Li, H. L.; Chen, L. L.; Liu, Y. Q.; Yang, X.; Shi, Y., Identification and characterization of a new geminivirus from soybean plants and determination of V2 as a pathogenicity factor and silencing suppressor. Bmc Plant Biol 2022, 22, (1).

5. Ayre, B. G., Membrane-transport systems for sucrose in relation to whole-plant carbon partitioning. Mol Plant 2011, 4, (3), 377–394.

6. Kumar, R.; Bishop, E.; Bridges, W. C.; Tharayil, N.; Sekhon, R. S., Sugar partitioning and source-sink interaction are key determinants of leaf senescence in maize. Plant Cell Environ 2019, 42, (9), 2597–2611.

7. Zhang, X. X.; Wang, M.; Wu, T. T.; Wu, C. X.; Jiang, B. J.; Guo, C. H.; Han, T. F., Physiological and molecular studies of staygreen caused by pod removal and seed injury in soybean. Crop J 2016, 4, (6), 435–443.

8. Wei, Z.; Guo, W.; Jiang, S.; Yan, D.; Shi, Y.; Wu, B.; Xin, X.; Chen, L.; Cai, Y.; Zhang, H.; Li, Y.; Huang, H.; Li, J.; Yan, F.; Zhang, C.; Hou, W.; Chen, J.; Sun, Z., Transcriptional profiling reveals a critical role of GmFT2a in soybean staygreen syndrome caused by the pest Riptortus pedestris. New Phytol 2023, 237, (5), 1876–1890.

9. Murray, M. G.; Thompson, W. F., Rapid isolation of high molecular weight plant DNA. Nucleic Acids Res 1980, 8, (19), 4321–5.

10. Fletcher, M.; Locker, H.; Mitchell, A.; Gopurenko, D., A revision of the genus Orosius Distant (Hemiptera: Cicadellidae) based on male genitalia and DNA barcoding. Aust Entomol 2017, 56, (2), 198–217.

11. Kumar, S.; Stecher, G.; Li, M.; Knyaz, C.; Tamura, K., MEGA X: Molecular evolutionary genetics analysis across computing platforms. Mol Biol Evol 2018, 35, (6), 1547–1549.

12. Song, N.; Cai, W. Z.; Li, H., Deep-level phylogeny of Cicadomorpha inferred from mitochondrial genomes sequenced by NGS. Sci Rep-Uk 2017, 7.

13. Hutton, E.; Grylls, N., Legume ‘little leaf ‘, a virus disease of subtropical pasture species. Aust J Agric Res 1956, 7, 85–97.

14. Helson, G., The transmission of witches broom virus disease of lucerne by the common brown leafhopper, Orosius argentatus (Evans). Biological Science 1951, 4, 115–124.

15. Harding, R.; Teakle, D., Mycoplasma-like organisms as causal agents of potato purple top wilt in Queensland. Aust J Agric Res 1985, 36, (443–449).

16. Pilkington, L. J.; Gurr, G. M.; Fletcher, M. J.; Nikandrow, A.; Elliott, E., Vector status of three leafhopper species for Australian lucerne yellows phytoplasma. Aust J Entomol 2004, 43, 366–373.

17. Thomas, J.; Bowyer, J., Properties of tobacco yellow dwarf and bean summer death viruses. Phytopathology 1980, 70, 214–217.

18. Horn, N. M.; Reddy, S. V.; Reddy, D. V. R., Virus-vector relationships of Chickpea chlorotic dwarf geminivirus and the leafhopper Orosius-Orientalis (Hemiptera, Cicadellidae). Ann Appl Biol 1994, 124, (3), 441–450.

